# A Critical-like Collective State Leads to Long-range Cell Communication in *Dictyostelium discoideum* Aggregation

**DOI:** 10.1101/086538

**Authors:** Giovanna De Palo, Darvin Yi, Robert G. Endres

## Abstract

The transition from single-cell to multicellular behavior is important in early development but rarely studied. The starvation-induced aggregation of the social amoeba *Dictyostelium discoideum* into a multicellular slug is known to result from single-cell chemotaxis towards emitted pulses of cyclic adenosine monophosphate (cAMP). However, how exactly do transient short-range chemical gradients lead to coherent collective movement at a macroscopic scale? Here, we developed a multiscale model verified by quantitative microscopy to describe wide-ranging behaviors from chemotaxis and excitability of individual cells to aggregation of thousands of cells. To better understand the mechanism of long-range cell-cell communication and hence aggregation, we analyzed cell-cell correlations, showing evidence of self-organization at the onset of aggregation (as opposed to following a leader cell). Surprisingly, cell collectives, despite their finite size, show features of criticality known from phase transitions in physical systems. By comparing wild-type and mutant cells with impaired aggregation, we found the longest cellcell communication distance in wild-type cells, suggesting that criticality provides an adaptive advantage and optimally sized aggregates for the dispersal of spores.

**Author Summary:** Cells are often coupled to each other in cell collectives, such as aggregates during early development, tissues in the developed organism, and tumors in disease. How do cells communicate over macroscopic distances much larger than the typical cell-cell distance to decide how they should behave? Here, we developed a multiscale model of social amoeba, spanning behavior from individuals to thousands of cells. We show that local cell-cell coupling via secreted chemicals may be tuned to a critical value, resulting in emergent long-range communication and heightened sensitivity. Hence, these aggregates are remarkably similar to bacterial biofilms and neuronal networks, all communicating in a pulse-like fashion. Similar organizing principles may also aid our understanding of the remarkable robustness in cancer development.

## Introduction

Many living systems exhibit collective behavior, leading to beautiful patterns found in nature. Collective behavior is most obvious in animal groups with clear advantages in terms of mating, protection, foraging, and other decision-making processes [60, 72]. However, how cells form collectives without visual cues is less well understood [41]. There are two main strategies to achieve synchrony (or long-range order) among individuals: A leader, i.e. a special cell or an external chemical field, may influence the behavior of the others in a hierarchical fashion (top-down). An example is the developing fruit-fly embryo in a maternally provided morphogen gradient [3, 45]. Alternatively, all individuals are equivalent and order emerges spontaneously by self-organization (bottom-up). Examples may include organoids [69] and other cell clusters [26], and both strategies are not mutually exclusive. While order itself cannot be used to differentiate between the two mechanisms, the response to perturbations or, simply, the correlations among fluctuations can be examined [4]. In top-down ordering, fluctuations are independent as cells follow the leader or the external field, and hence are not influenced by their neighbors. In contrast, in bottom-up ordering, cells are coupled to their neighbors. Hence, fluctuations are correlated as neighboring cells influence each other [10]. Note that in this context it is a reasonable assumption that cells can follow fluctuations of their neighbors much more easily than fluctuations of a distant leader cell. At a critical value of the cell-cell coupling strength, correlations may establish among cells which span the whole cell collective independent of its size, leading to a maximally connected collective similar to neurons in the brain [12].

To test these ideas of achieving order and long-range communication, we considered the well-known social amoeba *Dictyostelium discoideum*, which undergoes aggregation in response to starvation [8, 17, 35]. During this developmental process, cells start to secrete pulses of cAMP, a molecule that also acts as a chemoattractant for the other cells in the vicinity. The underlying signaling and regulatory pathways of such development have been thoroughly examined using genetics and imaging [19]: when a cell is ‘hit’ by a high concentration of cAMP, it secretes a pulse of cAMP itself, relaying the signal and thus causing the formation of cAMP waves, inferred indirectly from optical density waves in dark field movies [54, 73]. These waves propagate through the whole population [2, 21, 39, 62]. As their development proceeds, cells pulse at higher frequencies, reaching frequencies of up to one pulse every five minutes in the aggregate [20, 57]. Cell movement also accompanies the secretion process: before cells start to secrete cAMP, they normally move incoherently; when cAMP waves form, cells move towards the direction of the incoming wave by following the cells emitting the pulse in an orderly fashion (streaming phase). Interestingly, in a microfluidic device cells did not follow an artificially produced cAMP wave once it passed the cells, despite producing a gradient behind the cells pointing in the opposite direction of cell movement. Hence, cells are thought to solve the so-called ‘back-of-the-wave’ problem for directed unidirectional chemotaxis towards the aggregate [47, 59]. While singlecell chemotaxis [32, 42, 47, 52, 59, 66] and large-scale pattern formation [27, 33, 34, 38, 68, 73] have been extensively studied, a precise characterization of the transition from single cells to multicellularity is still missing.

Here, we developed a multiscale model to capture the mechanism of aggregation, focusing on the distinction between induced and self-organized order. Specifically, we were able to unify single-cell behavior and multicellularity at wide-ranging spatio-temporal scales. We achieved this by extending a single-cell model, which is able to describe *Dictyostelium* cell shape and behavior [66], by adding intracellular cAMP dynamics, secretion, and extracellular dynamics for cell-cell communication. To simulate hundreds of cells, we extracted a set of minimal rules for building a coarse-grained model. Hence, our approach is able to capture all stages of aggregation, ranging from single-cell chemotaxis to the multicellular collective. For quantifying the transition from disorder (pre-aggregate) to order (aggregate), we employed the mathematical concepts of spatial information and directional correlations. We found that the transition occurs during the streaming phase, which resembles a critical-like point known from phase transitions in physical systems as extracted by finite-size scaling. In physical systems phase transitions are characterized by an abrupt change in the macroscopic properties of the system when an external parameter (such as temperature) crosses a well-defined value. In our cell system, this parameter is the cell density (or external cAMP concentration). Criticality was tested by corresponding analyses of previously recorded time-lapse movies from fluorescence microscopy (provided by the Gregor lab [20, 49, 58]). Comparison of different *Dictyostelium* strains showed that wild-type cells have a longer cell-cell communication range than any mutant strain with impaired aggregation (based on regA and rdeA mutant data from the Cox lab [57]), even if cells with enhanced cell-cell adhesion (such as cells which secrete less cell number ‘counting factor’ [18]) form larger clusters. Hence, criticality may give cells an adaptive advantage, leading to optimally sized aggregates.

## Results

### A single-cell model fulfills criteria for aggregation

To model the transition from single cells to multicellularity, we started with cell shape and behavior in single cells. Specifically, we considered a model capturing single-cell membrane dynamics similar to the Meinhardt model [42, 48, 66]. Although not based on specific molecular species, this model describes membrane protrusions (such as pseudopods) and retractions, as well as resulting cell movement by means of three effective equations (see *Supporting Information)*. The first and second variables are a local activator and a global inhibitor (both are also considered in the local-excitation global-inhibition, LEGI, model [32, 52]). The third is a local inhibitor, important to destabilize the current pseudopod and to increase the responsiveness of the cell (Fig 1A, left). To this, we added equations representing the internal cAMP dynamics based on the FitzHugh-Nagumo model (Fig 1A, middle) [49, 58]. These are meant to capture the intracellular cAMP dynamics, governed by the relative activities of adenylyl cyclase ACA, which synthesizes cAMP, and 3’,5’-cyclic-nucleotide phosphodiesterase RegA, which degrades cAMP. Based on experimental evidence we assumed that cAMP is released from the rear of the cell [29, 40]. We also modeled extracellular cAMP dynamics for cell-cell communication, taking into account diffusion of cAMP in the extracellular medium and its degradation by secreted phosphodiesterase PDE (Fig 1A, right; see Materials and Methods for further information and *Supporting Information* for numerical implementation).

**Fig 1:**
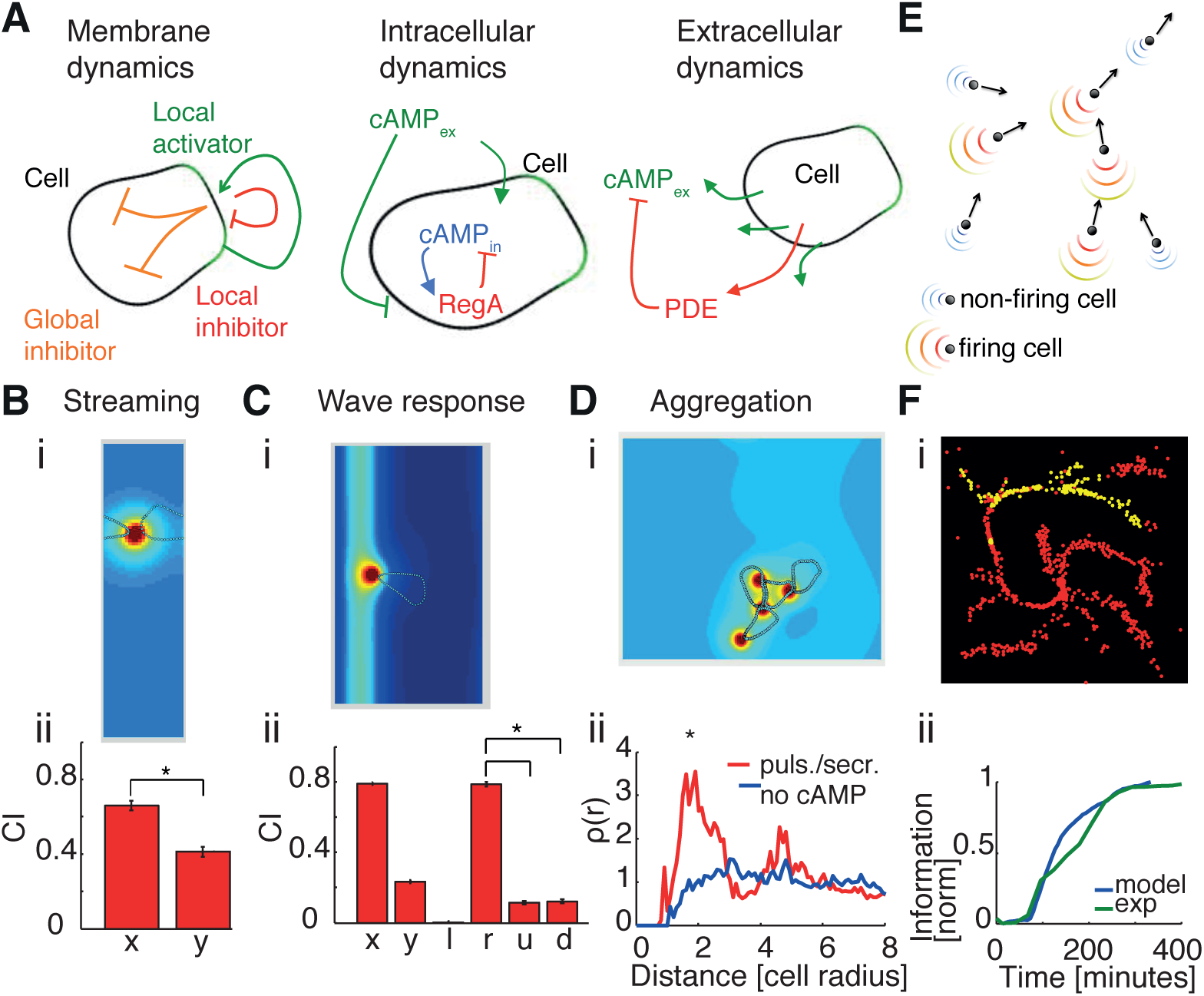
Multiscale model: from single cell-shape changes and chemotaxis to collective behavior. (A)Schematics for membrane dynamics (left), intracellular cAMP dynamics (center), and extracellular cAMP dynamics (right). (B) Single-cell “streaming” simulation in a box with periodic boundary conditions and a constant concentration of cAMP (i). Box dimensions are about 25×90 *μm* (the initial cell radius is assumed ~ *15μm*). Due to the small dimension of the box the cell is just leaking, not pulsing, in order to avoid saturation of secreted cAMP. The simulation was repeated 12 times and the average chemotactic index (CI) calculated (ii). Errorbars represent standard errors. Differences in CI_*x*_ and CI_*y*_ are statistically significant (p<0.01), using a Kolmogorov-Smirnov test (KS test). (C) Cells solve ‘back-of-the-wave’ problem. (i) A Gaussian wave (σ^2^ ~ 60*μm*) moves from right to left with a speed of about 300 *μ*m/min [67]. At the peak of the wave, the cell emits a pulse of cAMP. After the firing, the cell enters a refractory period during which it can neither fire again nor repolarize. The cell generally moves to the right, and hence does not follow the passing wave. (ii) CI in *x* and *y*, as well as in left (negative *x*), right (positive *x*), up (positive *y*) and down (negative *y*) directions, in order to discriminate between the directions of the incoming (right direction) and outgoing (left direction) wave. Simulations are repeated 12 times; shown are averages and standard errors. Box is about 60×105 *μm*. CI in the right direction is significantly higher than CI in the other directions. (D) “Aggregation simulation”. (i) Four cells are simulated moving in a constant concentration of cAMP. At the beginning, cells are randomly distributed. (ii) Density correlation at the end of simulations is plotted for control cells without secretion (blue) and all cells leaking cAMP and one cell also emitting pulses of cAMP (red). The red line has a significant (p<0.05, KS test) peak at a distance of about two cell radii, representing cell-cell contact. Also in this case simulations are repeated 12 times. Box dimensions are 75×75 *μm*. See Materials and Methods for details on density correlation, *Supporting Information* for a full explanation of the detailed model. (E) Schematic showing cells represented as point-like objects with velocity vectors. Firing cells emit pulses of cAMP, non-firing cells secrete cAMP at a low constant leakage rate. Spatial cAMP profiles are derived from detailed model simulations. At every time point cells are allowed two possible directions of movement in order to reproduce pseudopod formation at the cell front, with directions changing by ±27.5° with respect to the previous movement, corresponding to an angle between pseudopods of about 55° [70]. (F) Screenshot during streaming for *N* =1000 simulated cells (i). Red (yellow) points represent non-firing (firing) cells. (ii) Spatial information versus time: simulations (blue) compared with experimental dataset 3 (green). Values were then normalized and shifted in time to facilitate comparison.

Using this detailed model, we investigated the behavior resulting from cell-cell interactions in very small systems. First, we wanted our model to capture streaming, i.e. the ability of a cell to precisely follow the cell in front of it. To reproduce that, we simulated a single cell in a rectangular box with periodic boundary conditions (see Fig 1B and S1 Movie). In mathematics, periodic boundary conditions mean that the box is neighbored by identical copies of the box. Thus, in practice, if a cell passes through one side of the box, it reappears on the opposite side with the same velocity. This effect also applies to the molecules surrounding the cell. Now, given the rectangular shape of the box with the long side in the vertical direction and the short side in the horizontal direction, a horizontally moving cell can sense its own secretion (as the box is neighbored by identical copies of the box and hence copies of the cell and cAMP). Hence, the front of the cell is able to sense the secreted cAMP at the rear of the neighboring cell. In contrast, a vertically moving cell is too far away from its rear and thus cannot sense its secretion. We estimated the ability of the cell to stream by measuring the chemotactic index (CI) in the *x* direction, calculated as the amount of movement in the horizontal direction compared to the total length of the trajectory. In Fig 1B, we show that the CI in the *x* direction is significantly higher than the CI in the *y* direction.

We then considered the wave response as measured in microfluidic experiments, in which cells are exposed to traveling waves of cAMP [47, 59]. When hit by a traveling wave, cells moved towards the direction of the incoming wave but did not follow the wave after it passed. In order to capture this robust chemotaxis behavior, our model cell undergoes a refractory period (as was done in previous models [27, 33]) during which the cell can neither repolarize nor pulse (see *Supporting Information* for further details). Experimental evidence for this refractory period stems from the several minute-long directional bias in cell polarization [55], which may be caused by the large-scale inert cortical structure or phosphoinositide 3-kinase (PI3K), staying on the membrane even when no longer active [15]. In our simulations, this refractory state is naturally achieved when the cell spontaneously emits a pulse of cAMP upon encountering the wave (see Fig 1C, and S2 Movie). As a result, the CI is significantly higher in the right direction of the incoming wave. Finally, we considered a small number (four) of cells in a small box (with periodic boundary conditions) and tested whether they show signs of aggregation (see Fig 1D and S3 Movie). Specifically, we measured the tendency of the cells to cluster by calculating the density pair correlation function (see Materials and Methods), and compared the cases with and without secretion of cAMP. In the absence of secretion, cells were randomly distributed in space at the end of the simulations, as evident by the relatively flat horizontal line of unit correlation for distances larger than one cell length (the reduction at close distance is due to volume exclusion). With secretion, cells tended to be much closer to each other, with a clear peak in the density distribution at cell-cell contact (distance of two cell radii), indicating that cells tend to be close to each other and hence to cluster.

### A coarse-grained model reproduces collective behavior

In order to reproduce aggregation as observed in experiments, e.g. [20, 58], we need to simulate hundreds to thousands of cells. However, the detailed model introduced in the previous section is computationally too expensive, forcing us to introduce several simplifications. In our coarse-grained simulations, cells are point-like objects moving in continuous space. In particular, we took advantage of the spatio-temporal cAMP profiles from the detailed model by extracting the concentrations typically secreted by a single cell during leakage or a pulse. Shaped by degradation and diffusion, these profiles are approximately short-ranged exponential with a decaying amplitude in time. To capture the effects of volume exclusion, we also reduced the gradients in the cell-forward direction (see *Materials and Methods* for further information). Hence, as in the detailed model, the maximum cAMP concentration secreted by an individual cell is always found in the direction opposite to the direction of its motion. Using these analytical cAMP profiles, the cAMP concentration a cell senses is given by the sum of secretions by its neighboring cells. We then set concentration and gradient thresholds to determine whether a cell leaks or pulses cAMP, followed by a refractory period, and whether a cell moves randomly or follows the local cAMP gradient (see Fig 1E, Materials and Methods and *Supporting Information* for a detailed explanation of the model).

Using this minimal set of rules, we simulated thousands of cells with a density similar to the experimental ones (around a monolayer with 1 ML = 6, 600 cells/mm^2^ [20, 58]). Cells were initially distributed uniformly in space and allowed to move randomly. As soon as the cell density (and hence local cAMP concentration) increased spontaneously due to random cell motion, a cell may sense a concentration of cAMP large enough to pulse and this excitation will propagate throughout the whole population. Due to cell movement, streaming and aggregation into a small number of clusters can be observed (Fig 1F and S4 Movie). To quantify aggregation in a mathematical way, we estimated the ‘degree of order’ (or spatial information) in an image. This spatial information is based on the calculation of the 2D Shannon entropy, which does not require tracking of individual cells (see Material and Methods for mathematical details and *Supporting Information* for a primer on information theory) [24]. In this framework, evenly distributed cells correspond to a low spatial information, while highly clustered cells have a high spatial information. In all simulations, we observed that the spatial information rises sharply during the streaming phase as expected for cells in an ordered aggregate (see Fig 1F). Interestingly, the spatial information was previously used to capture the second-order (disorder-order) phase transition in the 2D Ising model (magnetic spins on a lattice) [24]. Hence, we wondered whether aggregation may be viewed as a critical-like point, describing the sudden transition from individual cells to the cell collective?

### Collective behavior: hierarchical or self-organized?

Based on our model assumptions, all cells are treated the same. However, aggregation may still be driven by the first random cell pulsing (hierarchical system) or spontaneously emerging as cells are coupled to each other by cAMP sensing and secretion (self-organized system; Fig 2A). The order of the collective process can be measured studying the directional correlations of pairs of cells. Specifically, the *non-connected (nc)* correlations

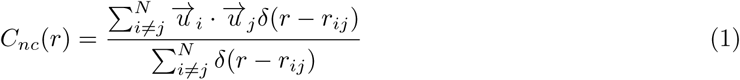

represent the average similarity of the direction of motion for every pair of cells depending on their distance, where N is the total number of cells, 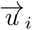 is the vector representing the direction of cell *i*, and 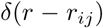 is equal to 1 if *r = r_ij_* and 0 otherwise. *C_nc_(r)* also represents the order parameter in our system, i.e. the quantity describing the degree of order or polarization in the system. For instance, when cells move independently of each other in random directions, then the order parameter is zero. In contrast, when all cells move in the same direction, then the order parameter would be maximal (i.e. one). By calculating this quantity for every time frame, we were able to analyze its variation in time. During the preaggregation stage correlations are close to zero even at short distances, while they increase sharply during the streaming phase (Fig 2B, top).

**Fig 2:**
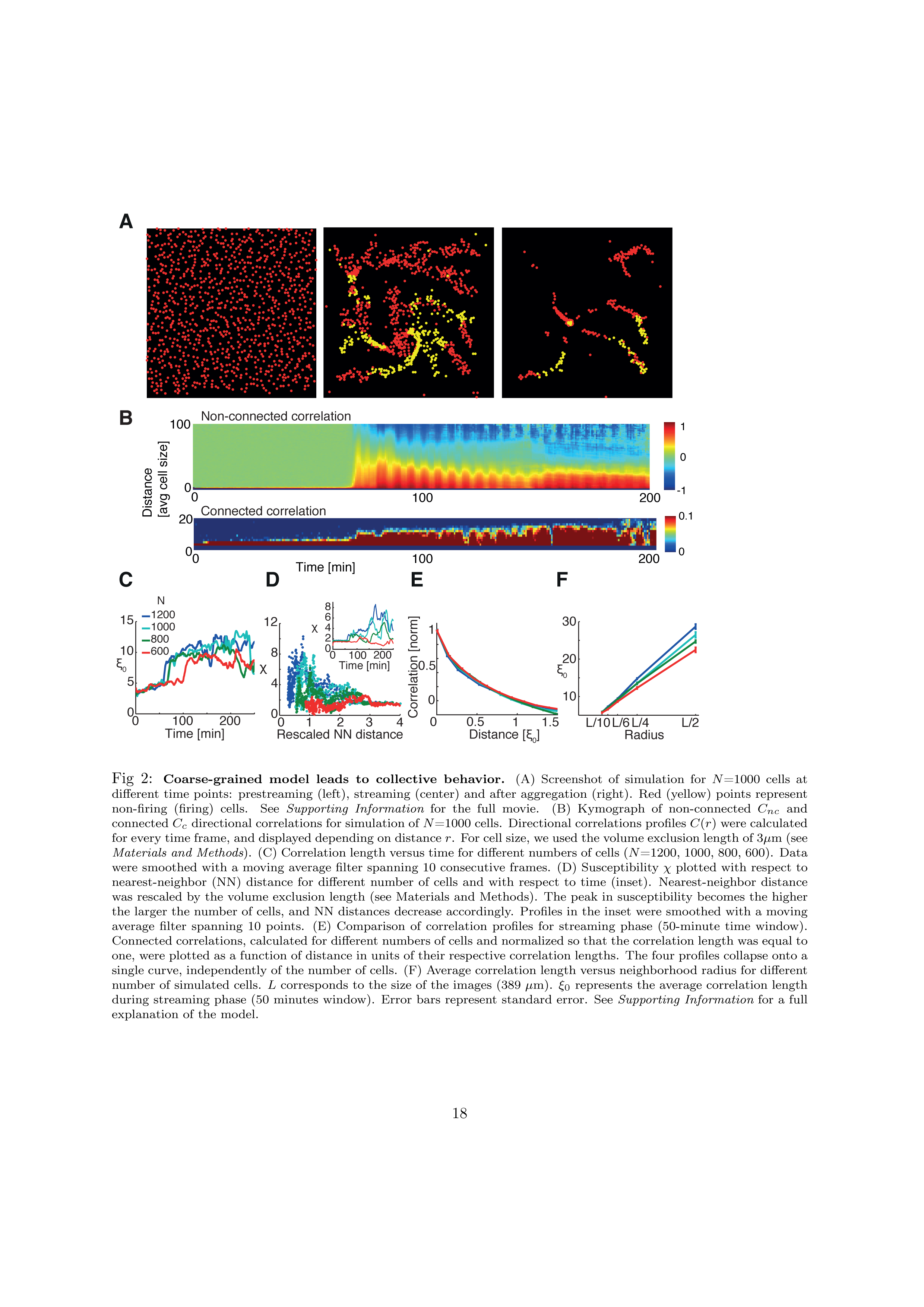
Coarse-grained model leads to collective behavior. (A) Screenshot of simulation for *N* =1000 cells at different time points: prestreaming (left), streaming (center) and after aggregation (right). Red (yellow) points represent non-firing (firing) cells. See *Supporting Information* for the full movie. (B) Kymograph of non-connected *C*_*nc*_ and connected *C_c_* directional correlations for simulation of *N* =1000 cells. Directional correlations profiles *C_(r)_* were calculated for every time frame, and displayed depending on distance *r*. For cell size, we used the volume exclusion length of 3μm (see *Materials and Methods*). (C) Correlation length versus time for different numbers of cells (*N* =1200, 1000, 800, 600). Data were smoothed with a moving average filter spanning 10 consecutive frames. (D) Susceptibility χ plotted with respect to nearest-neighbor (NN) distance for different number of cells and with respect to time (inset). Nearest-neighbor distance was rescaled by the volume exclusion length (see Materials and Methods). The peak in susceptibility becomes the higher the larger the number of cells, and NN distances decrease accordingly. Profiles in the inset were smoothed with a moving average filter spanning 10 points. (E) Comparison of correlation profiles for streaming phase (50-minute time window). Connected correlations, calculated for different numbers of cells and normalized so that the correlation length was equal to one, were plotted as a function of distance in units of their respective correlation lengths. The four profiles collapse onto a single curve, independently of the number of cells. (F) Average correlation length versus neighborhood radius for different number of simulated cells. *L* corresponds to the size of the images (389 μm). *ξ*_0_ represents the average correlation length during streaming phase (50 minutes window). Error bars represent standard error. See *Supporting Information* for a full explanation of the model.

Although order increases during the streaming phase, the origin and characteristics of this order are yet to be determined. To achieve this, we need to know more than the fact that the directions of cell movement are correlated (which describes the degree of order even in a hierarchical top-down system). In addition, we also need to know if the fluctuations of the directions are correlated. This would describe to what level cells communicate with each other and how they would respond collectively to perturbations.

For this purpose, we calculated the *connected* (c) directional correlations *C_c_(r)*, measuring the similarity of the directional fluctuations with respect to the average velocity [4, 10]. For instance, C_c_ = 0 (C_c_ = 1) means that a change in a cell’s direction is independent of (perfectly matched by) changes in the direction of its surrounding cells. To obtain the connected correlations, direction 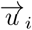 in Eq. 1 is substituted by the velocity of the single cell when the average is subtracted, i.e. 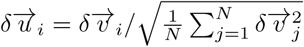 with 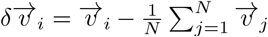. For this kind of collective movement, such a subtraction is not straightforward. If we compute a *global average* velocity for every time frame, we systematically overestimate the non-connected correlations, because we still consider part of the “bulk” velocity vectors due to the position of the cells in the image (see *Supporting Information* for a schematic explanation). To reduce this artefact, we considered *local averages*. For every cell, we considered the average velocity of all cells in its neighborhood up to a certain maximal distance *r_c_*, and we computed the correlations between the cell in the center and all the cells belonging to its neighborhood, and repeated this procedure for every cell in our image.

When applied to the simulations, Fig 2B, bottom, shows significant connected correlations, especially during streaming. Next, we considered the correlation length *ξ*_0_, i.e. a cell’s “influence radius” over its surrounding cells. We estimated this correlation length by the minimum distance at which the correlations cross zero, i.e. *C(r = *ξ*_0_)* = 0 [4]. We found that *ξ*_0_ is indeed much larger than the minimum nearest-neighbor distance. This indicates that a cell influences other cells way beyond its immediate neighbors, strongly suggesting self-organization (Fig 2C).

### Streaming as a critical-like point

Above, we demonstrated that aggregation in *Dictyostelium* is highly ordered and self-organized, with a correlation length much greater than the nearest-neighbor distance. Does the transition from disorder to order in this finite system show signs of criticality, given by a drastic and sudden qualitative change in behavior? At criticality all cells would remarkably influence each other independent of the distance between them.

In order to answer this question we considered that in critical systems the correlation length should scale with the size of the system as there is no intrinsic length scale [4]. To investigate this, we analyzed how the correlation length *ξ*_0_ changes in time. In all simulations *ξ*_0_ was small before aggregation and increased markedly during the streaming phase (Fig 2C). In equilibrium phase transitions, the susceptibility describes how sensitive the system is to perturbations, and this quantity would diverge at the critical point for an infinitely large system. Thus, this divergence indicates that the whole (infinite) system responds coherently as a single unit. In our cell system, the susceptibility can approximately be computed by the integrated correlations, i.e. by the amount of correlated cells,

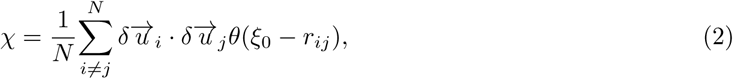
where *θ(ξ_0_ − r_ij_)* is equal to 1 for *r_ij_ < *ξ*_0_* and 0 otherwise [4]. This proxy for the susceptibility peaks precisely during the streaming phase (Fig 2D, inset), and the higher the number of cells, the higher the susceptibility. Moreover, if we consider cell density as a control parameter (similar to temperature or coupling in a ferromagnetic Ising model), we can plot χ with respect to the rescaled nearest-neighbor distance (Fig 2D and Materials and Methods). The resulting peak heights do not only reflect the number of cells, but their positions also shift to smaller nearest-neighbor distances (i.e. cells become more densely packed) as the number of cells increases, further supporting the resemblance to a scale-free system near criticality [4]. In theory, this peak height should keep increasing with cell number, and ultimately diverge at a critical nearest-neighbor distance for an infinite system. Furthermore, normalizing the correlations (so that they are one at the start for small distances) and rescaling the distance by the correlation length, the correlations collapse for all our simulations (Fig 2E). This collapse of the curves shows that they all have the same shape upon rescaling, indicating self-similarity as often occurring at criticality [25]. Finally, we took advantage of our image partition with different radii *r_c_* to examine how the correlation length *ξ*_0_ scales with system size. We noticed that for all movies, higher cell numbers display longer correlation lengths for a given neighborhood radius, and that the correlation length increases with increasing radius. Hence, the correlation length scales with system size (Fig 2F), indicating critical-like behavior in our simulated cells.

### Analysis of time-lapse fluorescent microscopy

To test the model, we analyzed five previously recorded movies of *Dictyostelium* aggregation with different cell densities from [20, 58] (see Materials and Methods and S5 Movie). Briefly, during 15 hours of observation, individual cells become a single, multicellular organism, going through different stages including preaggregation, streaming and aggregation (see Fig 3A). Cell densities ranged from 1/3 ML to almost 1 ML, ensuring aggregation while restricting our system to 2D. A 10% subpopulation of cells expressing the TRED fluorescent marker were tracked using a custom-written software (see Materials and Methods). Based on these cells, we repeated the analysis from the simulated cells for the TRED cells from the experiments, applying spatial information, non-connected and connected correlations, correlation length, and susceptibility. As we used the same computational protocol for both simulations and data, a close comparison was possible, which allowed us to assess finite-size scaling and hence critical-like behavior.

**Fig 3:**
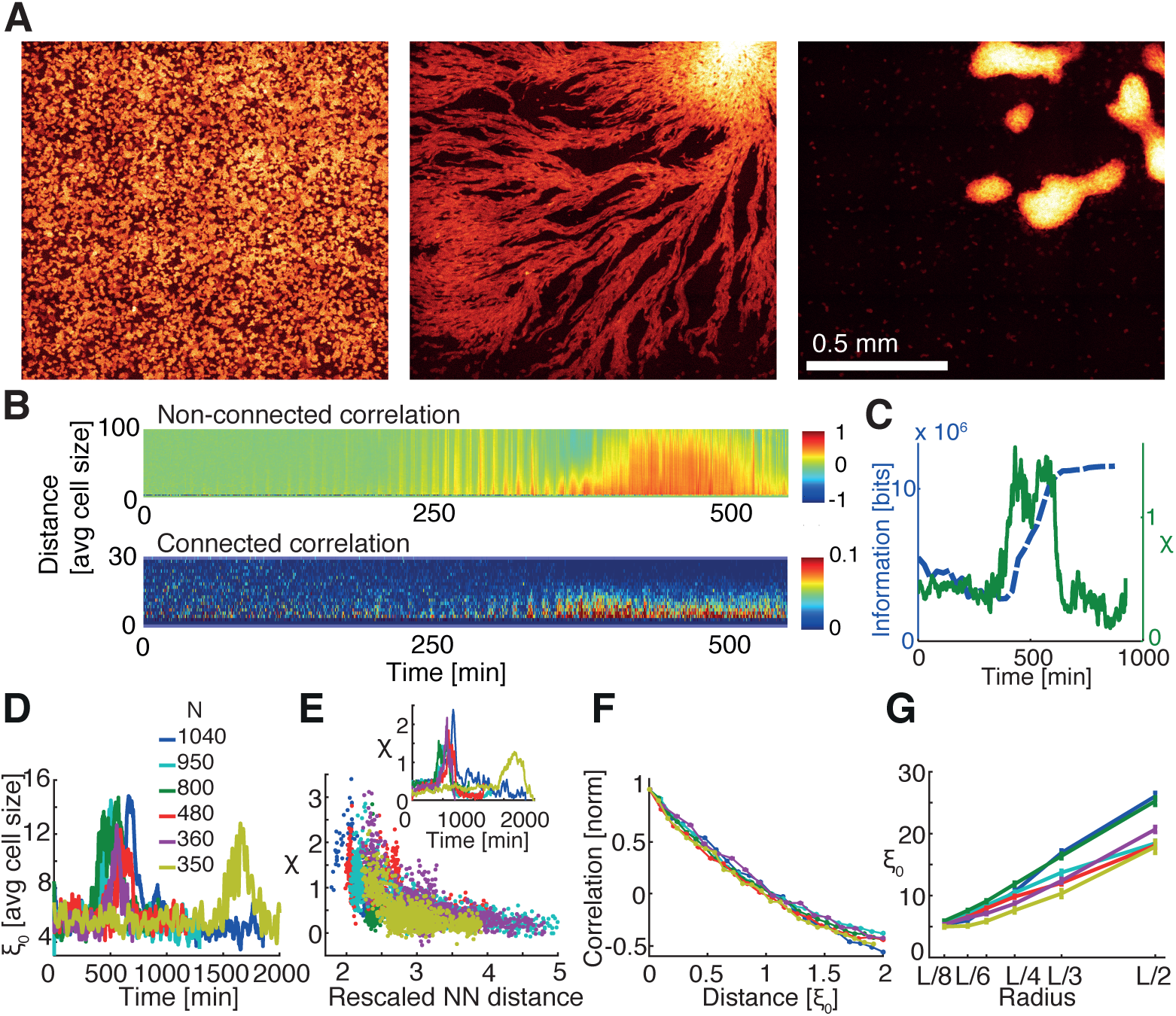
Collective behavior in experimental data for validating model. (A) CFP images of *Dictyostelium* aggregation of dataset 3. Images were taken after 4-5 hours of starvation when cells were still moving randomly before initiating aggregation (left), during streaming phase (450 minutes after first image, center) and after aggregation (800 min, right). (B) Kymograph of non-connected *C_nc_* and connected *C_c_* directional correlations for movie in dataset 3. Distance *r* is expressed in units of average cell size (estimated after an ellipse was fitted to every cell contour, and corresponding to the average of the minor axis, ~ 10.7*μm*) (C) Spatial information (blue) and susceptibility χ (green) of movie in dataset 3 as a function of time. The increase in spatial information denoting a more ordered image corresponds to the peak in susceptibility. (D) Correlation length *ξ*_0_ as a function of time for the six movies. Curves were smoothed with a moving average operation spanning 20 time points for better visualization. Inset: comparison of cell number estimated from TRED images during streaming phase for different movies. (E) Susceptibility χ as a function of rescaled nearest-neighbor (NN) distance and as a function of time (inset). Note that height of peaks increases and that the corresponding rescaled NN distance decreases with number of cells, as for simulations. Rescaled NN distance was computed by normalizing NN distance by the average cell size. In order to decrease noise, profiles in the inset were smoothed with a moving average spanning 20 time points. (F) Normalized *C_c_* as a function of correlation lengths *ξ*_0_ for different movies. *C_c_* for every dataset was calculated as an average over 150 minutes of streaming phase. Error bars represent standard error. As in the simulated data curve collapse for different numbers of cells. (G) Average correlation length versus neighborhood radius. *L* corresponds to the size of images (2033 pixels, ~ 1.3 mm). *ξ*_0_ represents the average of 150 min during streaming phase. Error bars represent standard error.

Based on our analysis of the data, the correlation length *ξ*_0_ increases during the streaming phase, as does the susceptibility χ (Fig 3B-E). Additionally, χ increases with cell number (and hence cell density), and the nearest-neighbor distance decreases, similar to the simulations. The correlation profiles, normalized and rescaled by the nearest-neighbor distance, largely superimpose for the different cell numbers, indicating that the slope of the resulting curves is not affected by the number of cells (see Fig 3F). Note, the cell density changes slightly over the duration of observation due to open boundary conditions (observation field is smaller than the field of cells so cells can freely move in and out of observation field; see *Supporting Information* for a quantification). Hence, cell numbers reported refer to the streaming phase (Fig 3D, inset). Finally, we studied how the correlation length changes for different system sizes by considering different neighborhood radii as performed for the simulations (see Materials and Methods). We noticed that *ξ*_0_ increases for a given radius with increasing cell numbers, and also for a fixed number of cells with increasing neighborhood radius (Fig 3G). These observations strongly suggest that there is no intrinsic correlation length, but that this length scales with system size. Taken together, our results suggest that aggregation can be viewed as a critical-like point in this finite system.

### Wild-type cells are better at long-distance cell communication than aggregation-impaired mutants cells

While criticality leads to long-range cell-cell communication, what is its biological function in aggregation and ultimate spore dispersal? This question can also be raised from the perspective of modeling: many published models achieve aggregation [27, 33, 51] (although assessing potential differences in the quality of aggregation is difficult in retrospect). If aggregation is readily achievable, what does criticality add to aggregation? There might be two ways to interpret this paradox: One is that all successful models are fine-tuned to achieve aggregation and this special point is again our critical-like point. Alternatively, simple aggregation is easy to achieve but aggregation of thousands of cells into a single aggregate (or very few aggregates) is difficult, and requires a diverging correlation length and hence exceptionally good long-range cell-cell communication. In this context, criticality may help making this process robust to variability and obstacles in nature as often microscopic details do not matter near a critical point [37, 63, 64].

To address this important question, we altered model parameters or reduced earlier assumptions. Specifically, we conducted aggregation simulations of 500 cells with (1) uniform (radially symmetric) secretion of cAMP (instead of secretion from the cell rear), (2) increased sensing noise (to address the naturally occurring cell-to-cell variability), (3) additional cell-cell adhesion (by the TgrA/C adhesion system during late aggregation [36]), which was not part of our original model, and (4) mutant cells with asynchronized cAMP secretion (similar to the regA mutant [20]). Experimentally, it was found that regA and rdeA with a diminished phosphorelay ability, as well as PDE mutants are able to aggregate, albeit into smaller clusters without streaming [1, 9, 11, 56, 61]. Furthermore, mutants with decreased ‘counting factor’ secretion (countin, cf45-1, cf50, or cf60) and hence increased cell-cell adhesion form larger cell clumps [9, 18, 50]. These cell types are implemented in our simulations as described in the Supplementary Information.

To quantify the range of cell-cell communication and the quality of aggregation, we considered the correlation length during the streaming stage and the spatial information of the final aggregate, respectively. Fig 4 A, C shows that these additional modified simulations exhibit decreased correlation lengths and spatial information as compared to our previous simulations (wild-type cells). Surprisingly, this even applies to the simulations with enhanced cell-cell adhesion, which produce broader aggregates as compared to wild-type cells. Thus, strong cell-cell adhesion leads to strong order, but apparently this does not allow for sufficient flexibility during the aggregation process. These findings are not in contradiction to earlier modeling, in which uniform secretion and adhesion allowed streaming to occur [16]; our results simply show that secretion from the cell rear further improves long-range cell communication and cell streaming, and that early adhesion can be detrimental to streaming. Subsequently, we estimated the correlation length and spatial information from previously published movies of wild-type cells, as well as regA and rdeA mutants [56, 57] (for details see *Supporting Information)*. Similar to our simulations, we found that also in experiments wild-type cells have a larger correlation length and spatial information than any of the analyzed mutants (Fig 4 B). Note that the simulations with ‘sensing noise’ and ‘asynchronous secretion’ can be made even noisier, which would reduce the correlation length and spatial information even further to match the data better.

**Fig 4:**
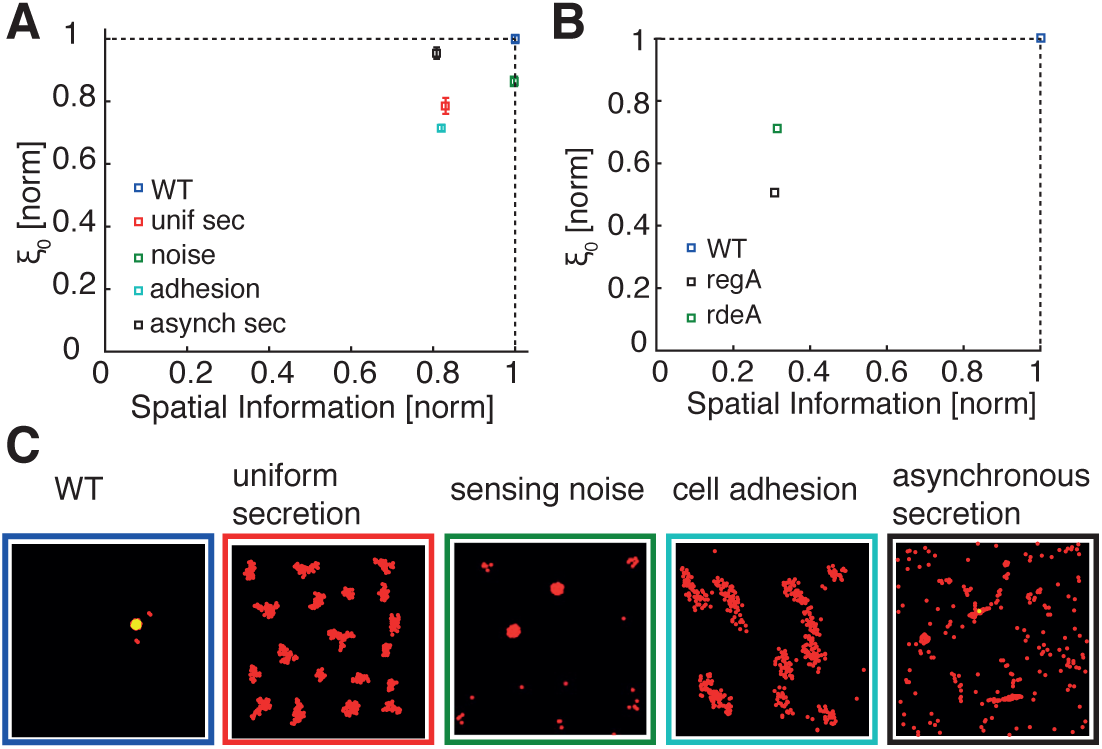
Role of criticality in aggregation of wild-type and mutant cells. (A) Correlation length (during streaming) and spatial information (of final aggregate) for coarse-grained simulations of *N* = 500 cells for wild-type (WT) and modified cell types. Correlation length and spatial information are normalized with respect to WT (blue symbol). Modified-cell simulations were performed with uniform (radially symmetric) secretion of cAMP (red), significantly increased sensing noise (tenfold increase in standard deviation compared to WT noise; green), enhanced cell-cell adhesion (light blue) and asynchronized secretion (random pulsing; black). (B) Corresponding correlation length and spatial information for experimental data from fig 5 of [56], considering WT cells (blue) and protein-kinase-A (PKA) pathway mutants (with the asynchronous regA mutant in black and the phosphorelay-intermediate-protein (rdeA) mutant in green). (C) Screenshots show cell distributions at the end of the simulations from (A). Error bars represent standard errors in correlation length for an average in time of 50 minutes during the streaming stage. See *Supporting Information* for a detailed explanation.

Hence, criticality may allow wild-type cells to create aggregates of just the right size, and, we speculate, may also be useful for decision making on where to aggregate, as well as for increased robustness in presence of obstacles and cell-to-cell variability. For instance, if aggregates are too small, stalks may be too short to disperse spores efficiently or too thin to support the weight of the spores [9]. In contrast, if aggregates are too large, then cells may encounter difficulties in decision making or cell sorting, or their stalks may collapse under the weight of too many spores. Hence, criticality may allow cells to make informed decisions for achieving optimal aggregate sizes for most effective spore dispersal. This would indicate that criticality constitutes an adaptive advantage.

## Discussion

*Dictyostelium* aggregation represents a fascinating example of synchronous collective cell behavior, spanning ~ 1mm in length although cells are just ~ 10*μ*m in size. Here, we asked how cells achieve such exquisite long-range communication [65], when the transition from single cells to the collective occurs, and how this transition can be characterized quantitatively. To capture the main features of aggregation, we developed a multi-scale model. First, we focused on single cells using a detailed model combining sensing, cell-shape changes and movement with cAMP secretion/pulsing and hence cell-cell communication. Once this model resembled the behavior of a single cell or a small group of cells, it allowed us to extract a minimal set of rules that could lead to aggregation. In particular, we extracted the cAMP concentration profile of a pulse from the detailed simulations and the refractory period after pulsing. By allowing cells to leak cAMP and to randomly move below a certain cAMP threshold concentration, we were able to observe spontaneous random pulsing as soon as the local density increased, similar to what occurs in real cells. This minimal set was subsequently included in the coarse-grained agent-based model, which is able to reproduce the collective behavior of hundreds of cells in line with time-lapse microscopy [20, 58].

Our major findings point towards previously uncharacterized features in aggregation, both observable in simulations and data. First, the transition to the collective is exactly pinpointed by a sharp rise in the spatial information of the cells during streaming. Second, to quantify the nature of the transition, we used fluctuations around the mean velocity, allowing us to distinguish between a hierarchically driven, top-down (external gradient from leader cells) and an emergent, self-organized, bottom-up (all cells are equal) process. Third, similar to second-order phase transitions in physical systems, the streaming phase shows signatures of criticality using finite-size scaling arguments. As a result there is no intrinsic length scale, allowing cells to communicate with each other over large distances ‘for free’, i.e. only based on local cell-cell coupling. The control parameter is cell density, affecting the cell-cell coupling via cAMP secretion and sensing.

Our work provides further insights into the process of cell aggregation. By means of our multi-scale model, we were able to answer why cells emit cAMP in pulses. Albeit short-lived, a pulse creates a steeper spatial cAMP gradient than continuous secretion (assuming that the total amount of emitted cAMP is the same in both cases). Moreover, we noticed that so-called cAMP ‘waves’ are likely not actual macroscopic traveling waves due to strong dissipation and diffusion. In contrast, cells are exposed to short-range cAMP pulses, which need to be relayed from one to the next cell before they dissipate. Although cAMP waves from microfluidic devices were used to study the cellular response to positive (incoming wave) and negative (passing wave) gradients, they may not represent natural stimuli [47, 59]. Hence, cells may not have to solve the traditional ‘back-of-the-wave’ problem, but instead have to decide which pulse to follow. However, this difficulty is eased as cells secrete cAMP from their rear [29]. Indeed, experiments of constitutively expressed adenylyl cyclase show defective streaming [30].

Our multi-scale model captures true emergence, generally not included in previous models of *Dictyostelium* aggregation. Models of wave propagation and spiral wave patterns go back as early as the 1970s [14], but generally these models did not include cell motility (but see [13] for an exception). More elaborate models from the 1990s focused on actual aggregation [27,33,34,39,51,73]. These were followed by the biologically more detailed LEGI [32, 52] and Meinhardt [42, 48] models to address the single-cell response to chemoattractant gradients. More recently, the FitzHugh-Nagumo model was adopted to explain the pulsing and synchronization of multiple cells (see *Supporting Information* for a comparison) [49,58], although early attempts to understand cAMP oscillations and the signal relay were already conducted in the 1980’s [39]. Furthermore, hybrid models were proposed [6]. However, none of these models started from a detailed spatio-temporal single-cell model and was able to quantify the cell-cell correlations, type of order, and exact transition point for achieving collective behavior.

When dealing with complex biological phenomena, there are necessarily limitations in the deduced models and acquired data. To assess criticality via finite-size scaling, ideally cell density is varied by orders of magnitude. However, this is often difficult to achieve in biological systems, and depends on experimental conditions. On the one hand, if cell density is much lower than about 1/3 ML, cells do not aggregate [58] (although lower density aggregation was achieved in a different experimental setup [23]). On the other hand, if the cell density is higher than 1 ML, experiments would need to be conducted in 3D with major technical difficulties. Despite the approximations, our model allows the identification of the key ingredients for certain observed behavior. For instance, an earlier version of our model showed some level of aggregation but no finite-size scaling. By investigating this shortcoming, we noticed that streams were too narrow due to nearly negligible volume exclusion. However, quasi-one dimensional streams restrict cell movement and suppress criticality, reminiscent of the missing disorder-order phase transition in the 1D Ising model according to the Mermin-Wagner theorem [43]. (Note that the 2D Ising model is a borderline case, but it is still possible to formally define a phase transition according to Kosterlitz and Thouless [28].) In our simulations, only when volume exclusion is increased and streams become broader, critical-like behavior emerges (see also discussion in [64]).

In an attempt to unify wide-ranging biological phenomena, short-range interactions may play similar roles in cell collectives (*Dictyostelium*, neurons, biofilms, embryos, tumors) [12, 31, 44] and animal groups (such as bird flocks) [4, 7, 10, 46, 71]. Interestingly, many different cell types communicate by pulsing (spiking), including neurons and bacteria [53]. Operating at criticality, i.e. the tipping point between order and disorder, may allow cells to be maximally responsive, to communicate robustly over long distances, to act as a single coherent unit, and to make decisions on, e.g., when and where to aggregate. In the future, it would be fascinating to conduct aggregation experiments in 3D environments, and to study the collective response to perturbations such as obstacles, changes in temperature, and exposure to toxins.

## Materials and Methods

### Detailed model

The intracellular cAMP dynamics are described by the FitzHugh-Nagumo model, a classical model to reproduce neuronal spiking and previously adopted to describe excitability in *Dictyostelium* [49, 58]. Degradation of intracellular cAMP is achieved by phosphodiesterase RegA, which is negatively regulated by extracellular concentration of cAMP (by means of extracellular signal-regulated kinase ERK2 [40]). Secretion of cAMP from the cell rear [29, 40] is strictly coupled to its intracellular concentration: if the extracellular cAMP concentration is below a threshold value cells exhibit a constant small leakage of cAMP, but a temporary high concentration of cAMP is released during pulses of intracellular cAMP once above the threshold. If the extracellular cAMP concentration is kept above this threshold the cell becomes a sustained oscillator. Extracellular cAMP is degraded by the phosphodiesterase PDE [5]. This model correctly captures the relay of the signal and the sustained pulsing observed in *Dictyostelium* (see *Supporting Information* for a detailed explanation).

### Coarse-grained model

To reproduce the dynamics of thousands of cells, we simplified further the representation given by the detailed model. We assumed that cells are point-like objects, which secrete cAMP maximally at their rear. Specifically, spatial propagation of cAMP was modeled as an exponential decay with a constant of 0.1 *μm*^−1^ (within a factor of 2 of the value extracted from the detailed model simulations). The spatio-temporal concentration profiles are rescaled according to the cosine of the angle with the opposite-to-motion direction; secretion becomes zero at 90° (lateral secretion) and is set to zero for all the frontal part of the cell. (The above mentioned fine tuning of the exponential decay constant may be a result of this rescaling approximation, or may reflect the fact that the cell-cell coupling is a key parameter for critical-like behavior.) We set a concentration threshold *c*_1_ to determine if a given cell will emit a pulse or just leak cAMP, and a gradient threshold ∇*c*_2_ determines if the cell will move randomly or follow the local cAMP gradient. As for the detailed model, every cell undergoes a refractory period of 6 minutes after firing, during which it keeps the same motion it had during pulsing. To reproduce volume exclusion, cells cannot be closer to each other than 3*μm* (this rule is overwritten later in simulations, when cells are densely packed and likely superimpose). To drastically speed up simulations, the algorithm is written without explicit modeling of diffusion of cAMP in space, instead it computes how much cAMP every cell senses and what their spatial gradients are by considering positions of cells with respect to each other. This implementation is able to reproduce aggregation of thousands of cells. More specifically, *N* =1000 cells were considered at experimental density of about one monolayer (1ML=6, 600 cells/mm^2^). For the other simulations of *N* =600, 800 and 1200, the total area (of 389×389 μm) was fixed and density varied accordingly. See *Supporting Information* for a detailed explanation.

### Density pair correlation

The pair-correlation function was computed as described in [22], given by

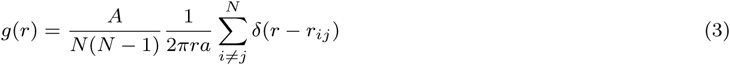
where A is the total area of the image considered, N is the number of cells, r is the radius of a ring and *a* is the discretization constant. In case of a random distribution *g(r)* takes a value of 1 on average (similar to blue trace in Fig 1Dii), while in case of particle clustering *g(r)* becomes greater for small distances (as for red trace in same panel).

### Spatial information

Spatial information of an image of cells was calculated in Fourier space of wave numbers based on the formalism described in [24]. All images were binarized (by means of MATLAB thresholding algorithms *graythresh* and *im2bw* for the case of experimental images). After that, 2D images were converted in 3D binary matrices where the third dimension has a 1 corresponding to the pixel intensity (thus in this case, since the starting images were binary, the 3D matrix has a 1 at level 0 if that pixel is black and at level 1 if it is white). This guaranteed that all images had the same histogram, provided that they initially were of the same size. For the case of uncorrelated pixels, all Fourier coefficients *P_i_* are considered independent and Gaussian distributed. Image entropies were then calculated as:

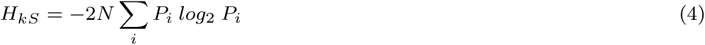
where the probability density function *P* is Gaussian distributed with zero mean and variance calculated from the sum of the pixel intensities. *H*_*ks*_ is computed by dividing the function into bins of width σ/100 and summing *P*_*i*_ *log*_2_ *P*_*i*_ from − 10σ to 10σ. Fourier transformation was then applied to the image. The real and imaginary part of the Fourier coefficients were then considered to compute

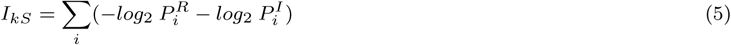
where 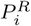 and 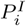 refers to the real and imaginary part of coefficient *i*. The sum was calculated by considering bins of width σ/100 around the values assumed by the Fourier coefficients. k-space spatial information *kSI* was finally calculated as *kSI* = *H*_*ks*_ − *I*_*kS*_. For a primer on information theory, see *Supplementary Information*.

### Directional correlations and susceptibility

To calculate the connected correlations, local averages of the velocities were subtracted from cell velocities. For every cell we considered the average movement of all cells in its neighborhood up to a certain maximal distance *r*_*c*_, and compute the correlations between the cell in the center and all the cells belonging to its neighborhood. We repeated this procedure for every cell in our image. In this way we are able to decrease the “bulk” velocity component in the fluctuations, while keeping a continuous partition of the image (which we would have lost in case of rigid partition of the image in smaller squares), and without preassigning the final position of the aggregation center. In order to understand better the influence of this partitioning on the calculation of the connected correlations, we repeated the same procedure for different radii. Specifically, if *L* is the image dimension, we set *r*_*c*_ equal to *L*/2, *L*/4, *L*/6, *L*/8, and *L*/10, with *L*/6 appearing to be the best choice in terms of the trade-off between avoiding overestimation of correlations and number of cells in the neighborhood for good statistics in the simulated data. For the analysis of experimental data, *L*/2, *L*/3, *L*/4, *L*/5, *L*/6, and *L*/8, were considered, and *L*/4 was chosen, reflecting again the trade-off between good statistics of noisy dataset and small overestimation of correlations. To plot the susceptibility, we estimated the nearest-neighbor distance, computed for every frame as the average of the nearest-neighbor distances for all cells.

### Experimental methods

Time-lapse movies were obtained similar to protocol in [20, 58]. Axenic *Dictyostelium* cells expressing the Epac1camps FRET sensor were starved for 4-5 hours, and then plated on hydrophobic agar for imaging. Sixteen fields of view from a microscope are combined (1.2 × 1.2mm^2^), resulting in the recording of thousands of cells in a wide field (inverted epifluorescence microscope (TE300, Nikon). To allow high-precision tracking of individual cells in a dense cell population, a different fluorescent marker, mREPmars (TRED), was expressed and mixed with unmarked cells so a subpopulation of cells could be tracked (10% TRED cells). See *Supporting Information* for further details.

### Segmentation and tracking

Images of TRED channels were segmented by using the MATLAB function *imextendedmax*, which outputs a binary image given by the computation of the local maxima of the input image. The centroids positions were then computed from this mask by means of the *regionprops* function. The tracking of individual cells was done by considering the centroid positions for different times. For every time *t* the nearest neighbor centroid at time *t* +1 was found, and the trajectory was accepted if the distance between the two positions was smaller than the average cell size.

## Acknowledgments

We are grateful to the Gregor lab at Princeton University for sharing their data with us, and additionally Thomas Gregor, Allyson Sgro, and Monika Skoge for helpful discussions and comments on the manuscript. We also thank Luke Tweedy for help with the detailed model, Javier López-Garrido and Linus Schumacher for a critical reading of the manuscript, and Mariam Elgabry and Suhail Islam for support with computational issues. GDP and RGE were supported by ERC Starting Grant 280492-PPHPI (http://erc.europa.eu/starting-grants).

